# Immune Profiling among Colorectal Cancer Subtypes using Dependent Mixture Models

**DOI:** 10.1101/2023.07.24.550400

**Authors:** Yunshan Duan, Shuai Guo, Wenyi Wang, Peter Mueller

**Affiliations:** Department of Statistics and Data Science, University of Texas at Austin; The University of Texas MD Anderson Cancer Center

**Author notes:** Corresponding author: Peter Mueller.

## Abstract

Comparison of transcriptomic data across different conditions is of interest in many biomedical studies. In this paper, we consider comparative immune cell profiling for early-onset (EO) versus late-onset (LO) colorectal cancer (CRC). EOCRC, diagnosed between ages 18-45, is a rising public health concern that needs to be urgently addressed. However, its etiology remains to be poorly understood. We work towards filling this gap by identifying homogeneous T cell subpopulations that show significantly distinct characteristics across the two tumor types, and to identify others that are shared between EOCRC and LOCRC. Such inference may reveal underlying determinants of clinically observed differences in the two disease subpopulations. We develop dependent finite mixture models where immune subtypes enriched under a specific condition are characterized by terms in the mixture model with common atoms but distinct weights across conditions, whereas common subtypes are characterized by sharing both atoms and relative weights. The proposed model defines a variation of mixtures of finite mixture models, facilitating the desired comparison by introducing highly structured multi-layer Dirichlet priors. The model allows us to explicitly compare features across conditions. We illustrate inference with simulation studies and data examples. Results identify EO-enriched and LO-enriched T cells subtypes whose biomarkers are found to be linked to mechanisms of tumor progression. The findings reveal distinct characteristics of the immune profiles in EOCRC and LOCRC, and potentially motivate insights into treatment and management of CRC.

## 1 Introduction

Early-onset colorectal cancer (EOCRC) refers to CRC diagnosis in individuals aged 18-45 (or ≤ 50) and is an increasing global health problem that needs to be urgently addressed [1]. An important gap in current research is that the differences in immune cell populations of EOCRC and late-onset colorectal cancer (LOCRC) have not been well investigated [2]. We develop inference for immune profiling of EOCRC and LOCRC samples, aiming to identify, compare, and understand relations of immune cell subtypes across these two disease subpopulations. The problem is formalized as statistical inference in two dependent mixture models, where immune subtypes are represented as components in the mixture. The specific aim of identifying differences in immune subtype abundance between the two conditions is achieved by introducing a highly structured multi-level prior across the two linked models. Under each disease condition, the mixture model gives rise to a partition, i.e., a cluster arrangement of observed immune cells. Inference can thus alternatively be characterized as joint clustering of all immune cells under the two conditions, using synchronized random partitions.

We proceed under a model-based Bayesian paradigm. Bayesian inference for a mixture model Σ_*h*_ *w*_*h*_*f* (*y* | *θ*_*h*_) is often based on representing the model as a mixture of a kernel *f* (*y* | *θ*), e.g., a Gaussian kernel, with respect to a discrete mixing measure 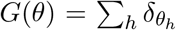 (here *δ*_*x*_ is point mass at *x*). The construction is motivated by Kingman’s representation theorem [3] which shows that under certain symmetry conditions (exchangeability) any random partition can be thought of as arising this way. A prior probability model for the random mixing measure *G* completes the model construction. Priors for random distributions are known as nonparametric Bayes (BNP) models [4], and the by far most widely used BNP model is the Dirichlet process (DP) prior [4, Chapter 4]. We write *G ∼* DP(*α, G*_0_), where *α >* 0 is a precision parameter and *G*_0_ is the prior expectation of the random probability measure. One important property of the DP prior is the a.s. discrete nature of *G*, i.e., 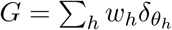. The DP provides a computationally attractive model for an unknown probability measure *G*. However, for inference on two related random mixtures, as needed in our motivating application, we need a prior probability model *p*(*G*_1_, *G*_2_) for two dependent random probability measures *G*_1_ and *G*_2_, characterizing the immune subtypes under the two disease conditions. We build on an extensive literature on such models over the past 20 years. Some of the earliest constructions are additive decomposition models introduced in [5], which use a mixture of common components and idiosyncratic components, both defined by discrete random probability measures with DP priors. The strength of the dependence is controlled by the relative weight of common versus idiosyncratic parts. Kolossiatis et al. [6] introduced a restriction that ensures that the marginal prior on *G*_*j*_ remains a DP. Lijoi et al. [7] generalized such models using random probability measures constructed via completely random measures (CRM).

Other dependent models *p*(*G*_*j*_; *j* = 1, …, *J*) that are built on DP priors include the hierarchical and the nested DP and variations thereof. Teh et al. [8] introduce the hierarchical DP (HDP) which constructs a probability model for dependent random measures *G*_*j*_ by linking submodels *G*_*j*_ *∼* DP(*M*_*j*_, *G*) with a common base measure *G*, which in turn is given a DP(*M*_0_, *G*_0_) hyperprior. In contrast to the HDP, the nested DP (NDP) proposed by Rodriguez et al. [9] links the random probability measures *G*_*j*_ by assuming *G*_*j*_ *∼ Q*, using a common prior 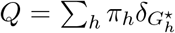. Here *Q* is a distribution of distributions. A sample *G*_j_ from *Q* selects one of the atoms 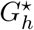with probability *π*_*h*_. Assuming a hyperprior *Q ∼* DP(*α*, DP(*M, H*)) creates a discrete distribution *Q* with each atom 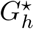 itself being generated by a DP, 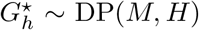. The latter implies marginally a DP prior *G*_*j*_ *∼* DP(*M, H*). We shall later discuss and compare more properties of HDP and NDP, as they are relevant for our motivating application. A generalization of the HDP is studied in [10–12] by replacing the DP with more general normalized completely random measures (NRM). A completely random measure (CRM) is a random measure *μ* with the defining property that *X* = *μ*(*A*) and *Y* = *μ*(*B*) for two mutually exclusive events *A ∩ B* = *∅* are independent random variables [13]. The hierarchical NRM includes the HDP and the hierarchical Pitman-Yor (PY) process, another variation with the same construction, as special cases. Camerlenghi et al. [14] extend the NDP by defining the latent nested process (LNP) as a nested process of NRM’s, which is constructed by normalizing an additive structure including shared and idiosyncratic CRM components. More recently, Denti et al. [15] proposed a nested common atoms model (CAM), where different distributions are characterized by specific weights assigned to a common set of atoms. Yet another construction that combines elements of the HDP and NDP models is the hidden hierarchical Dirichlet process (HHDP) model of Lijoi et al. [16]. Like in the NDP they generate the *G*_*j*_’s from a prior *Q*. But the atoms 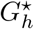of *Q* are generated from a HDP, as discrete random probability measures with shared atoms. The HHDP blends features of the HDP and the NDP. Additionally, instead of assuming fully exchangeable data in the models above, models for partially exchangeable data have been investigated using nested clustering [17], and separate exchangeable structure [18].

Considering the immune profile comparison problem, differential immune cell behaviors are expected when comparing EO-with LOCRC, i.e., some immune subtypes are shared across conditions while some occupy a significantly larger proportion in one condition than the other. Suitable inference models should therefore include shared immune subtypes with common weights and common atoms across all conditions, and idiosyncratic but not exclusive components with shared atoms, but distinct weights. The previously summarized constructions do not introduce and favor such structure, making them less suitable for the application in comparative immune profiling. One notable limitation is that the idiosyncratic components that appear in some of the constructions are independent, lacking clusters with common atoms but distinct weights. The HDP and related priors, as well as the CAM and HHDP model do not allow for components with common weights and atoms. While the CAM and the HHDP model include common atoms across all conditions, there is no structure in the model to favor a distinction into shared clusters versus clusters that are enriched under different conditions. Finally, it is natural to assume a finite but unknown number of immune subtypes, advising against the use of infinite mixture models.

The discussed limitations prompt us to consider alternatives. We build on finite mixture models. Recent discussions of Bayesian inference for mixtures of finite mixtures (MFM) appear in Frühwirth-Schnatter et al. [19], Greve et al. [20] and Miller and Harrison [21]. These constructions create a single random probability model *G* as a finite mixture of random size *K*. MFM models appear to be more suitable than DP models in many examples and offer many of the same features. For example, when focusing on the implied clustering of experimental units, the probability function of the implied random partition is characterized by exchangeable partition probability functions (EPPF) with similar properties as under corresponding BNP mixtures. A natural extension of MFM to be suitable for the motivating application involves multiple, related mixtures.

In this paper, we introduce a structured dependent finite mixture model, including model features that anticipate the desired inference in the comparative immune profiling problem. We construct the model using an additive structure of common components and components that are specific to each condition. The model is useful for any inference problem concerned with clustering of features and comparison across different conditions.

The rest of this paper is organized as follows. Section 2 describes the motivating immune profiling problem and the data sources. We describe explanatory data analysis and preprocessing steps for data cleaning and dimension reduction. In Section 3.1, we introduce a dependent finite mixture model (“888 model”) for a general inference problem. In Section 3.2 we introduce a specific implementation for the motivating application on immune profile comparison between EO and LO-CRC. We discuss the differences and commonalities with alternative models in Section 3.3. More properties and details of posterior inference are included in Section 4.1 and 4.2. In Section 5, we discuss simulation studies to validate finite sample performance of the model. Section 6 shows the results for the motivating comparative immune profiling problem. The findings in immune subtypes, especially the ones that are enriched under a certain condition reveal some underlying biological paradigm for the disease subtype.

## 2 Immune profiling of EOCRC vs. LOCRC

Recent studies have identified distinctive clinical features in EOCRC patients aged 18-45 (or ≤ 50) compared to LOCRC patients over 60 [1]. Also, it is known that the immune cells within the tumor microenvironment (TME) profoundly influence tumor development, progression, and response to treatment in CRC patients [22, 23]. This leads us to hypothesize that the immune microenvironment of EOCRC differs from that of LOCRC and promotes tumor progression. To further investigate the distinct immune microenvironment in patients with EOCRC, we develop a full probabilistic description of comparative immune profiling between EO and LO conditions. Such inference will help reveal some of the underlying determinants for the distinct etiology of EOCRC and for the potentially different treatment response between the two disease conditions.

Our analysis is based on single-cell RNA sequencing data obtained from Zhang et al. (2018) [22], which was sequenced using the Smart-seq2 platform (see **Appendix A**). This study only sequenced T cells, allowing us to focus on investigating the subtypes of T cells. Because CRC samples with microsatellite instability (MSI) have distinct immune profiles compared to microsatellite stable (MSS) CRC samples and are known to be more responsive to checkpoint inhibitor (CPI) treatment, we further removed tumor samples with MSI [24]. We followed the standard analyzing pipeline of the *Seurat* package for data preprocessing. Only genes that were detected in a minimum of three cells were retained. Next, we excluded low-quality cells by setting thresholds for the number of expressed genes and detected Unique Molecular Identifier (UMI) counts: cells were retained if they expressed between 500 and 6,500 genes and had UMI counts ranging from 500 to 2,000,000. Taking into account the information provided by the original paper [22], our study focused on CD4+ and CD8+ cells in tumors and adjacent normal tissues, while other T cell types (such as double-negative and double-positive cells) and cells originating from the peripheral blood were filtered out. After these quality control steps, we were left with a total of 3,139 CD4+ and CD8+ cells from seven CRC patients, including two EO and five LO patients. For data visualization we used Uniform Manifold Approximation and Projection (UMAP) transformations implemented with the *Seurat* package, as shown in Figure 1 (a)-(b). In this figure, panel (a) organizes the data by disease condition (EO vs. LO), and panel (b) does so by the type of cell (CD4+ vs. CD8+).

**Figure 1:**
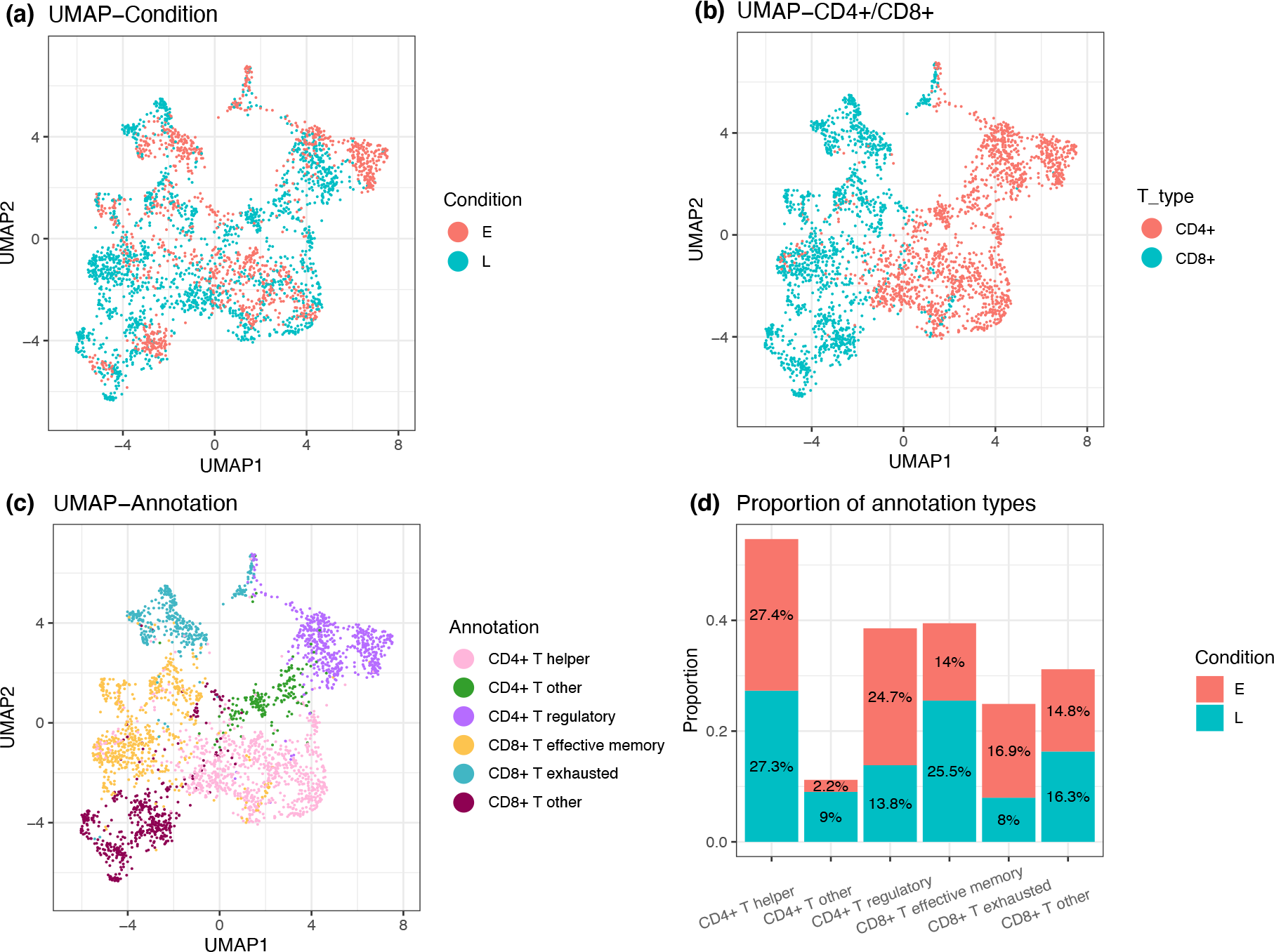
Single-cell RNA-seq data from CRC samples in UMAP plots, colored by EO/LO conditions (panel a), T cell CD4+/CD8+ types (panel b), and cell type annotations (panel c). Panel (d) shows the proportions of annotated cell types under EO and LO condition.

As part of exploratory data analysis, we first carried out clustering as implemented in the *Seurat* package, setting the resolution at 0.75. We annotated T cells into immune subtypes, as illustrated in Figure 1 (c). The annotation is based on matching markers for the known T-cell sub populations. CD4+ cells were annotated into three subtypes: regulatory T cells, helper T cells, and other CD4+ T cells. Similarly, CD8+ T cells were annotated into three subtypes: effective memory T cells, exhausted T cells, and other CD8+ T cells. The relative proportions of EO and LO conditions within each immune cell subtype cluster are depicted in Figure 1 (d). The results suggest a potential correlation between cell subtypes and onset conditions, which was formally tested using Fisher’s exact test. We observed a highly significant correlation (p=0.0005), indicating that the data contains sufficient information for the desired comparative immune profiling. However, an omnibus test of independence does not provide information about which subtypes are shared or characteristic for each condition. We also implemented k-Means clustering [25] (results not shown), which, however, also stops short of detecting interpretable immune cell sub-populations for the comparison between EO and LO conditions. In summary, these preliminary analyses indicate the feasibility of inferring disparities in immune cell sub-populations between EO and LO conditions and demonstrate the need for an inference approach specifically aiming at identifying comparative immune subtypes between conditions.

## 3 Dependent finite mixture model

### 3.1 888 model

We construct a dependent finite mixture model for comparative immune cell profiling to identify homogeneous immune cell sub-populations. Let *y*_*ji*_ denote gene expression features observed for cell *i* = 1, …, *N*_*j*_, under condition *j* = 1, …, *J* . We develop the model for a more general case with *J >* 1 conditions, keeping in mind that we have *J* = 2 for the motivating application. The proposed probability model is defined as follows. The features *y*_*ji*_ are generated from finite mixtures *f* (*·*;*θ*) *dG*_*j*_(*θ*) with respect to a finite, discrete mixing measure *G*_*j*_, *j* = 1, …, *J*, using, for example, a normal kernel ∫ *f* (*·*; *θ*). Writing the mixture as an equivalent hierarchical model with latent variables *θ*_*ji*_ we have:

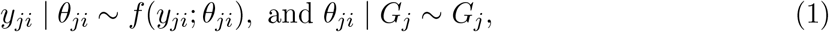

*i* = 1, …, *N*_*j*_ and *j* = 1, …, *J* . The desired dependence across *j* is induced by a hierarchical prior on *G*_*j*_

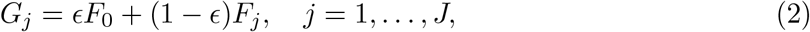

In words, the probability measure *G*_*j*_ for subjects under condition *j* is decomposed into a shared component *F*_0_ and condition-specific components *F*_*j*_. In anticipation of the comparative immune profiling problem, the latter is constructed to include some parts that are interpretable as sub-populations enriched under condition *j*, and others that are interpretable as enriched under conditions *ℓ* ≠*j*. This is implemented by representing *F*_*j*_ as a mixture of distributions 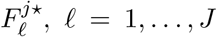, including distributions that characterize the subtypes that are enriched under condition *j* itself 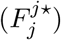, and in other conditions 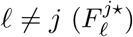, respectively:

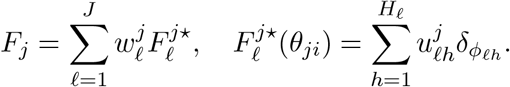

The 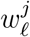 define the relative weights of components enriched under different conditions, and the 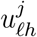 define the relative weights of different sub-populations that are specific to condition *ℓ*. A prior on the relative weights 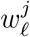 leads to this desired interpretation by imposing that (*a priori*) 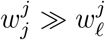, *ℓ* ≠ *j*. That is, all conditions share the same subtypes with the same atoms (cluster-specific parameters *ϕ*_*lh*_) but at different levels of relative abundance 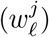. *A priori* under condition *j*, components in 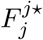 have a significantly higher abundance than components in 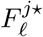, (*ℓ* ≠ *j*). Similar to the 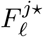, the shared measure *F*_0_ is also assumed to be discrete with finitely many atoms:

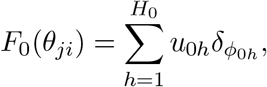

now with common (relative) weights, shared across all conditions, *j* = 1, …, *J* . Together the weights *ϵ*, 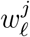, and 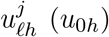 define a prior on the set of weights under conditional *j* for all unique cell sub-populations. The purpose of the hierarchical split into the three sets of weights is to allow the prior to favor a structure that anticipates the desired interpretations. In other words, the purpose of the structured hierarchical prior on the weights is prior shrinkage towards an interpretable structure with the split into sub-populations enriched under one or the other condition etc. However, such structure is only favored, not enforced. It is possible that *a posteriori* some of the idiosyncratic clusters are estimated with practically equal weight under different conditions, making them effectively shared clusters.

The mixture model induces a random partition of cells. Introducing latent cluster membership indicators *c*_*ji*_ and *s*_*ji*_ the model can alternatively be written as follows. First, there is a mixture of *f* (*y*_*ji*_; *ϕ*_*lh*_) with respect to 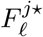 characterized by *s*_*ji*_:

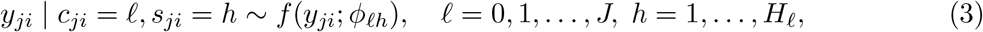

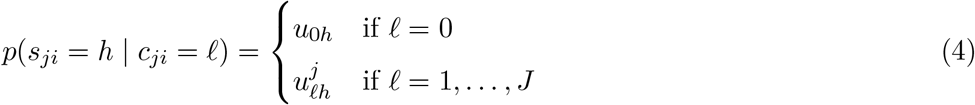

representing the mixture with respect to the *H*_*R*_ components of 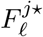 (when *c*_*ji*_ = *l >* 0), and with respect to the *H*_0_ components of *F*_0_ (if *c*_*ji*_ = 0). Another mixture occurs with the choice of the type of component *ℓ* in

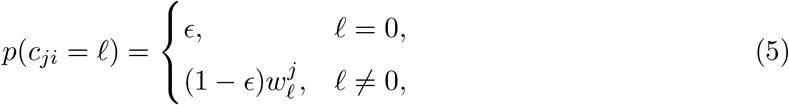

where *c*_*ji*_ is the indicator for *y*_*ji*_ arising from a common cluster (*ℓ* = 0) or an idiosyncratic cluster (*ℓ* ≠ 0). A third level of mixing could arise with possibly random *H*_*ℓ*_ and *H*_0_.

To ensure the structure of the different terms in the mixture, we introduce an asymmetric Dirichlet prior on 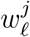:

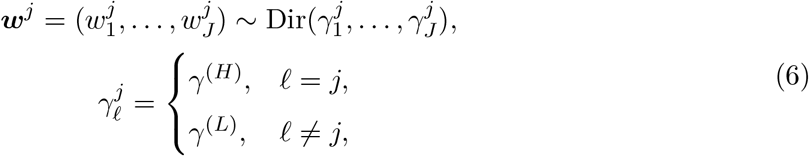

with *γ*^(*H*)^ ≫ *γ*^(*L*)^, allowing *a priori* 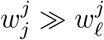, *ℓ* ≠*j*. A detailed explanation on how the prior model shrinks the abundance of clusters and introduces the desired structure for our inference goal appears in the specific example of immune profiling, and is included in Section 3.2. We assume symmetric Dirichlet and beta priors on other weight parameters.

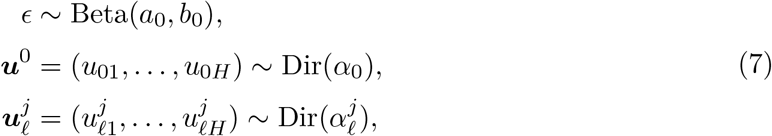

where Dir(*·*) with one parameter denotes a symmetric Dirichlet prior. Together (6) and (7) define for each condition j a prior on the set of weights for all unique atoms. The prior is defined hierarchically to allow the desired structure across conditions.

We refer to the model as “888 model”, keeping in mind the case with 2 conditions and 8 clusters in each of the three cluster types, i.e., *H*_0_ = *H*_1_ = *H*_2_ = 8. Note that the number of clusters *H*_*j*_’s can be random, or any fixed number in the general “888 model”, despite the choice of the name.

### 3.2 Comparative immune profiling for EO-versus LOCRC

We discuss specific model details in the implementation for the motivating application to immune profiling. For easier notation and without loss of generality, assume *H*_0_ = *H*_*ℓ*_ = *H*, and consider the *J* = 2 conditions as in the CRC immune subtypes example, with *j* = 1, 2 denoting the LOCRC and EOCRC condition, respectively. The dependent mixture model can be expressed in the following form,

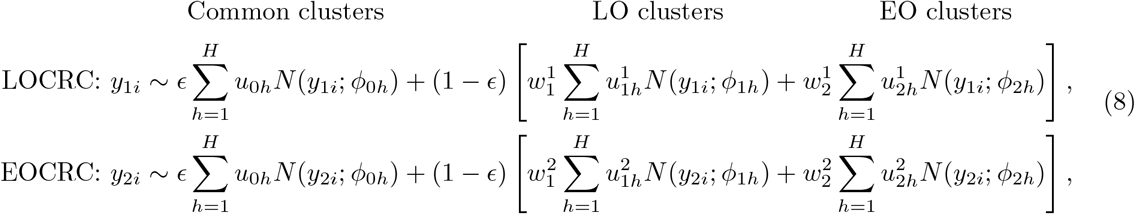

where *ϕ*_*ℓh*_ = *{ξ*_*ℓh*_, Σ_*ℓh*_*}, l* = 0, 1, 2, are the cluster-specific parameters, i.e., the mean and covariance matrix of the Gaussian distributions. The first *H* clusters are the common immune subtypes, the second set of *H* clusters are the LO-enriched subtypes, and the last *H* clusters are the EO-enriched subtypes. The atoms (cluster-specific parameters) *ϕ*_0*h*_, and *ϕ*_*ℓh*_ are shared between EOCRC and LOCRC conditions. Thus, the immune subtypes identified by the proposed mixture model share the same gene expression profile between EO and LO conditions, for both common and idiosyncratic clusters. The shared atoms in the idiosyncratic components in *F*_*j*_ are critical to allow for the desired comparison of immune cell profiles across conditions.

We assume that the common immune subtypes share common weights *ϵu*_0*h*_, while condition-specific clusters are characterized by distinct weights 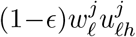, with *a priori* 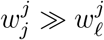, *ℓ* ≠ *j*, given by the prior. The EO/LO-enriched immune subtypes are prominent in samples of the matching condition, but not exclusively. In other words, EO-enriched clusters tend to have high weights in the EOCRC immune cell populations and low weights that are shrunk towards 0 in LOCRC immune cells, and vice versa for LO-enriched clusters.

To achieve such structure of comparative clusters in the model, an informative asymmetric beta prior is put on the idiosyncratic cluster weights 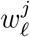for LO condition (*j* = 1) and EO (*j* = 2) condition, 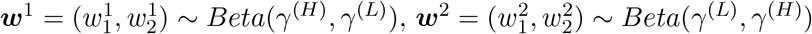, which is a special case of equation (6) when *J* = 2. We use a small value *γ*^(*L*)^ for the beta parameters corresponding to the cluster weights to be shrunk to 0, and a large *γ*^(*H*)^ for those weights to be shrunk away from 0. Symmetric priors for other relative weights, *u*_0*h*_, *u*_*lh*_, and *ϵ*, are specified in equation (7).

We use *γ*^(*H*)^ = 10, *γ*^(*L*)^ = 1, 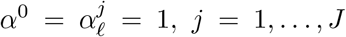, *a*_0_ = 10, and *b*_0_ = 10, in the simulation studies and data analysis, which serves as suitable prior hyperparameters for the desired structure. For example, under the EOCRC condition, the weights of LO-enriched clusters *w*^2^ are shrunk to 0, and the sum of weights in EO-enriched clusters is close to 1. This is consistent with our prior assumption of the immune subtypes that we are interested in.

For other parameters in the model, we assume conditionally conjugate priors on the cluster-specific parameters, 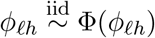. We use 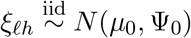 and 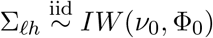 for *l* = 1, 2, *h* = 1, …, *H* in our examples. One could add a hyperprior on *H* if desired. In the application we choose to proceed with fixed *H*, to represent strong investigator preference for a moderate interpretable number of clusters for each condition.

### 3.3 Alternative models

Several similar inference approaches for dependent random probability models have been proposed in the recent literature, including the references that were already briefly cited in the introduction. We discuss some more details, including limitations that are relevant for the inference on immune cell subtypes.

Some alternative models for dependent random probability measures build on an additive structure where the discrete random measure *G*_*j*_ for condition *j* is a weighted mixture of DPs. The probability model for *G*_*j*_ is composed of a common component *H*_0_ and an idiosyncratic random measure *H*_*j*_,

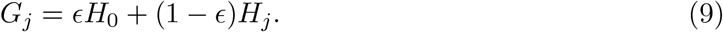

The model is completed with independent DP priors for *H*_*j*_, *j* = 0, 1, …, *J* . Such models were considered in [5] and further developed in [6] by restricting the prior probability model for *ϵ* to ensure marginal DP priors for *G*_*j*_, and in [7] by assuming that *H*_0_ and *H*_*j*_ arise by normalizing compeletly random measures (CRMs), including the DP as a special case. The common characteristics of these additive structure models is that the models across conditions share a common discrete measure *H*_0_ implying shared atoms and proportional weights for the common components across all conditions, as desired for the motivating application. However, the idiosyncratic components are entirely different, with no notion of shared atoms. This is not consistent with the structure of homogeneous immune cell sub-populations that are shared across conditions, even if they are characteristic and enriched under one or the other condition.

The hierarchical DP (HDP) model proposed by Teh et al. [8] defines the random probability measures *G*_*j*_ as DP random measures, with a shared base measure *G*, which in turn is generated by another DP hyperprior:

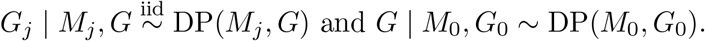

Camerlenghi et al. [10] generalize the HDP to hierarchical normalized random measures (NRM), with *G*_*j*_ and *G* defined as NRMs,

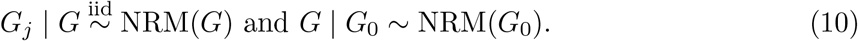

including the HDP as a special case with a particular choice of the CRM. This type of hierarchical construction generates discrete random measures with all shared atoms across conditions as in our proposed 888 model. However, although all atoms are shared across *G*_*j*_, actual allocated clusters, i.e., atoms that are (latently) linked with data, need not be shared. Also, none of the components in the model share weights, and the models with exchangeable components lack the structure of the weights that identifies the common and condition-specific immune subtypes.

The nested DP (NDP) proposed by Rodriguez et al. [9] assumes the random measures *G*_*j*_ are samples from a distribution *Q* of distributions, that is 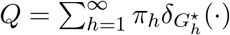 with the weights generated by stick breaking with total mass parameter *α*, and the 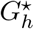 being generated by a DP prior, 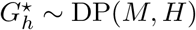. In short,

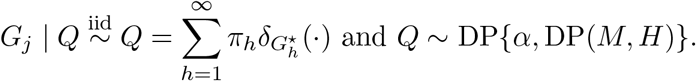

The nested process (NP) introduced in Camerlenghi et al. [14] constructed the same nested structure with CRM’s. Both NDP and NP generate random measures *G*_*j*_ that are either identical, sharing all the weights and atoms, or with all distinct weights and atoms. If *θ*_*ji*_ generated from *G*_*j*_ share at least one value across two conditions *j*_1_ ≠ *j*_2_, then the posterior degenerates on the case 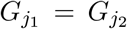. This would be inappropriate for an application with common immune subtypes across all conditions. Recognizing this posterior degeneracy, Camerlenghi et al. [14] introduced the latent nested process (LNP), which is an additive structure, composed of a common component from a CRM, and an idiosyncratic component from a NP. This model tackles the degeneration problem of NDP and NP mentioned above, and allows for some shared clusters across conditions. But the idiosyncratic components can only be either identical or have all distinct weights and atoms. The model does not allow for clusters with common atoms and distinct weights, and therefore fails to accommodate inference for differential abundance of shared sub-populations.

The hidden hierarchical Dirichlet process (HHDP) model introduced by Lijoi et al. [16] builds on the NDP structure, but assumes an HDP prior for the 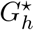 . Replacing independent sampling from DP(*M, H*) with non-atomic *H* by the HDP prior allows for shared atoms across different 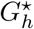.

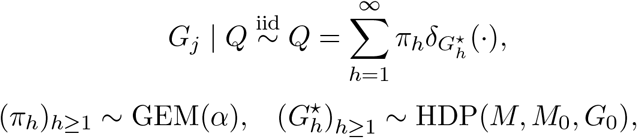

Here GEM stands for the distribution of probability weights after Griffiths, Engen, and McCloskey, which characterizes the stick-breaking construction in the DP prior. The distinct distributions 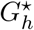 are dependent and have a common base measure leading to shared atoms across 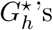. It overcomes the degeneracy problem of the NDP but introduces no structure to distinguish the clusters by the characteristics of the weights in HHDP, so that the clusters could not be identified as common or specific to a certain condition by the construction of the model.

## 4 Properties of the 888 model

### 4.1 Random partition probability function and large sample properties

The mixture model in equation (1) induces a random partition of the experimental units – in our case, the immune cells. Given the latent cluster assignment indicator 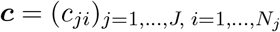 for common (*c*_*ji*_ = 0), or condition *ℓ* enriched clusters (*c*_*ji*_ = *ℓ* ≠ 0), the probability function of the random partition for the 888 model is characterized by a partially exchangeable partition probability function (pEPPF) [10]. The pEPPF is a function of cluster sizes that is symmetric with respect to all clusters for each of the cluster types defined by ***c*** and satisfies a coherence condition across sample sizes. To derive the probability function, note that 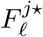 and *F*_0_ are the discrete random mixing measures in an MFM. Let Dir_*K*_(*γ*) denote a symmetric *K−*dimensional Dirichlet Dir(*γ*, …, *γ*) distribution. Consider a generic MFM model,

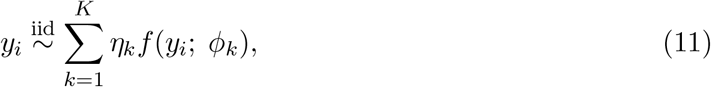

*i* = 1, …, *N*, with *p*(*η*_1_, …, *η*_*K*_) = Dir_*K*_(*γ*) and a prior *p*(*K*) on the size of the mixture. Let 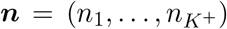 denote the cardinalities of *K*^+^ *≤ K* clusters implied by the ties of *N* = Σ_*k*_ *n*_*k*_ i.i.d. samples from (11). The random partition of the *N* experimental units (for example, cells), is characterized by the EPPF [19, 21],

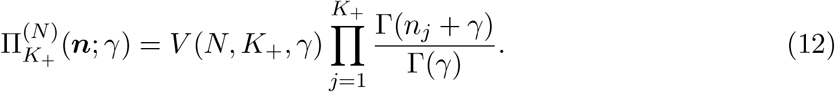

where

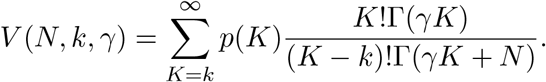

The random partition probability function for the 888 model can be derived based on (12) for each of the implied MFM models, and marginalizing out the *ϵ* and *w* weight parameters. We need some notation. For given data, let 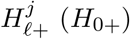 denote the number of clusters created by ties of sampling latent parameters from 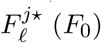. Let 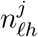 be the number of units (cells, in our case) from condition *j* allocated to the *h*th cluster enriched under condition *ℓ, j* = 1, …, *J, ℓ* = 1, …, *J*, 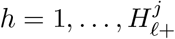, and 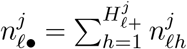 be the total number of units from condition *j* allocated to clusters enriched under condition *ℓ*. Similarly, let *n*_0*h*_ denote the number of units from all conditions being allocated to the *h−*th cluster of the common clusters, *h* = 1, …, *H*_0+_, and let *n*_0•_ be the total number of units allocated to common clusters.

#### Proposition 1

*The pEPPF given the cluster type indicator* ***c*** *is*

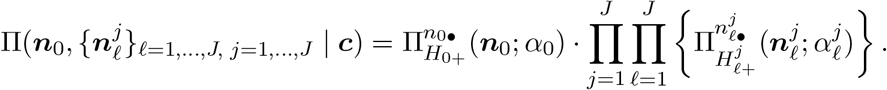

*The random partition probability function for the 888 model is*

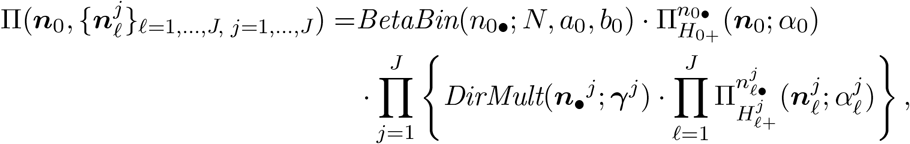

*where* 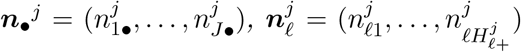, 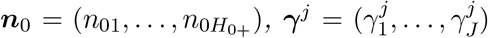. 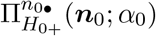 *and* 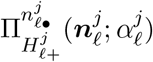 *are the EPPFs of the static MFM defined in equation* (12). *BetaBin*(*·*), *DirMult*(*·*) *denote the probability functions of a beta-binomial and Dirichlet-multinomial distribution, respectively*.

*Proof:* See the appendix.

An alternative representation of the random partition is given by the predictive probability function [26]. Let 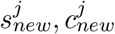 denote latent cluster membership indicators for a future *i* = (*N*_*j*_ + 1)*−*st sample from *G* . The predictive distribution for 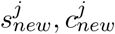 given 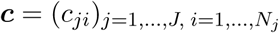 and 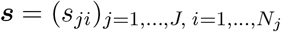 takes a form similar to the Pólya-urn under a DP prior [27]. The following expression is useful for constructing posterior MCMC transition probabilities.

#### Proposition 2

*The Pólya-urn representation of the predictive probability function under the 888 model is*

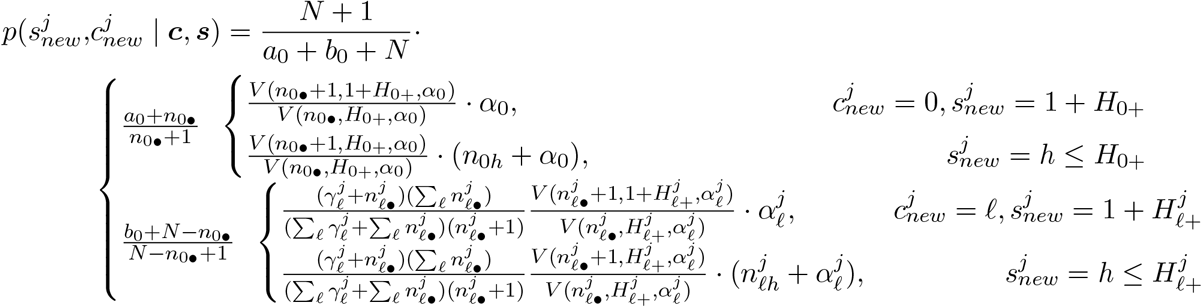

*where* 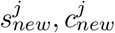 *denote the allocation of a new object in condition j, indicating the cluster and the type of cluster, respectively, as defined in equation* (3).

*Proof:* See the appendix.

To study large sample properties of posterior inference under the 888 model it is convenient to focus on the implied marginal model for condition *j*. The marginal model for *y*_*ji*_, *i* = 1, …, *N*_*j*_, can be written as a simple finite mixture model with all *J × H* clusters, and an asymmetric Dirichlet prior on the weights by using the aggregation property of the Dirichlet distribution, if assuming the following constraints on the prior model:

#### Lemma 1

*Assume the hyperparameters in the priors of the weights satisfy*

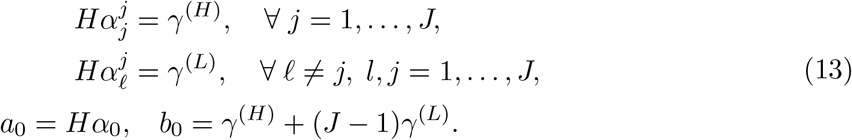

*Then the marginal model for y*_*j*_ *under the 888 model is equivalent to a MFM model with asymmetric Dirichlet prior:*

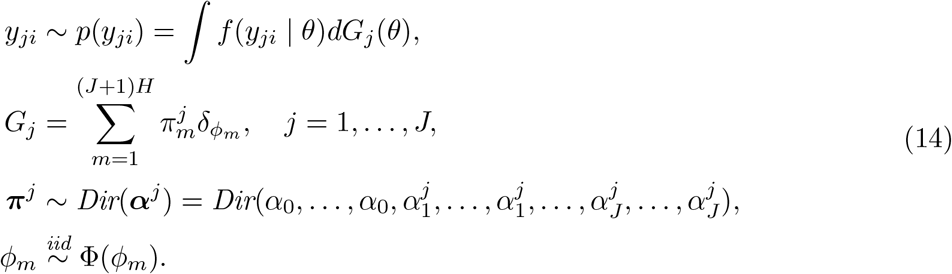

*Proof:* The claim is implied by the aggregation property of the Dirichlet distribution [28, Chapter 49].

In this case, the 888 model can be considered as a variation of the MFM model with an asymmetric Dirichlet prior that incorporates the desired structure.

We briefly discuss large sample properties of the posterior for the mixing measure *G*_*j*_. Assume that data *y*_*ji*_ are generated from a mixture with true mixing measure 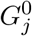. We show that for any condition *j*, the posterior distribution, 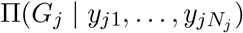 converges with increasing *N*_*j*_ to the true mixing measure 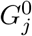, and the posterior contraction rate is given as follows.

#### Proposition 3

*Under model* (14), *for ∀j* = 1, …, *J, we have*

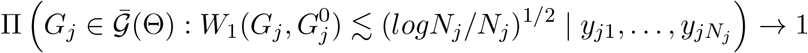

*in probability under* 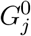. *Here*, 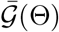 *denotes the space of all discrete measures, and W*_1_ *denotes the L*_1_ *Wasserstein distance*.

*Proof:* We give an outline of the proof. The claim follows from Theorem 3.1 in Guha et al. [29] which gives posterior contraction rates for a general MFM. Replacing the symmetric Dirichlet prior with the Dirichlet prior in equation (14), the condition on the prior probability of the parameters in a neighborhood of the truth used in the proof still holds. The assumptions (P.1) - (P.4) in [29] hold under our proposed model. Therefore, we can prove the posterior contraction rate following the scheme of proof in Guha et al. [29]. See the Appendix for details.

If we consider the finite case where the number of clusters *H < ∞* is fixed, the posterior distribution of the mixing measure can also be proven to be consistent, and obtain a posterior contraction rate of (log *N*_*j*_*/N*_*j*_)^1*/*2^. This follows from Guha et al. [29] and Ho and Nguyen [30].

In summary, for large enough sample size (under each condition *j*) inference under the proposed model can recover the underlying sub-populations close to the truth. Therefore, under large enough sample sizes in all conditions, inference on the comparison between conditions recovers the truth.

### 4.2 Posterior inference

In our implementation (in the simulation studies and the data analysis) we use mixtures with fixed *H*. This choice is motivated by the fact that the number of immune subtypes that we are interested in for biological interpretation is limited. The model is sufficiently flexible as long as *H* is chosen as an upper bound beyond the number of immune subtypes of interest. We summarize posterior inference by first reporting a point estimate of the partition by minimizing the variation of information (VI) loss as proposed in Dahl et al. [31]. To maintain the structure of the model, the search for the optimal partition is restricted to the posterior Monte Carlo samples of imputed partitions. The *salso* R package is used to calculate VI loss. Minimizing VI loss, we obtain an optimal cluster assignment *ĉ*_*i*_, and estimated cluster specific parameters 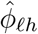, including the cluster centroids 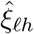 and the covariance matrices 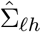.

We report uncertainty on cluster allocation of the cells given the cluster specific parameters. Considering the posterior mean of the log cluster assignment probability, we define a metric *q*_*ji*_(*ℓ, h*) as the similarity of cell *i* under condition *j* to cluster *h* among the type *ℓ* clusters,

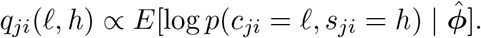

For a given cell, the higher the similarity to cluster *h*, the higher the probability that it is assigned to cluster *h*. Here 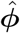 is a point estimate of the cluster specific parameters.

## 5 Simulation Study

We carry out a simulation study to evaluate the performance of the 888 model under realistic finite sample sizes and with single-cell sequencing data. The simulation shows how the model successfully identifies the true clusters and provides the desired inference on comparative abundances of cell type sub-populations between two conditions.

The simulated data is generated from a publically available dataset of Peripheral Blood Mononuclear Cells (PBMC) collected on a 10X platform [32]. The cell types given in the dataset are used as hypothetical simulation truth, and are denoted as types 1-4 in Figure 2 (a). We assume the cells are collected under two conditions A and B. We generate cell-specific conditions with a defined true probability *P* (cell *i ∈* condition B | cell *i* is of type *k*) = *p*_*k*_. In our simulation setting, *p*_1_ = 0.01, *p*_2_ = *p*_3_ = 0.99, *p*_4_ = 0.52. Therefore, under the simulation truth, cell type 1 is dominant in condition A, cell types 2 and 3 are enriched under condition B, and cell type 4 shares a similar, but slightly different proportion between two conditions. Figure 2 (b) plots the cells color coded by simulated (true) conditions using UMAP scores. Figure 2 (c) shows the proportion of the four true cell types under condition A and B, respectively. In the analysis we assume *H* = 2, that is, we assume *a priori* that the first two clusters are shared, the second two clusters are specific to conditions B, and the last two clusters are specific to conditions A.

**Figure 2:**
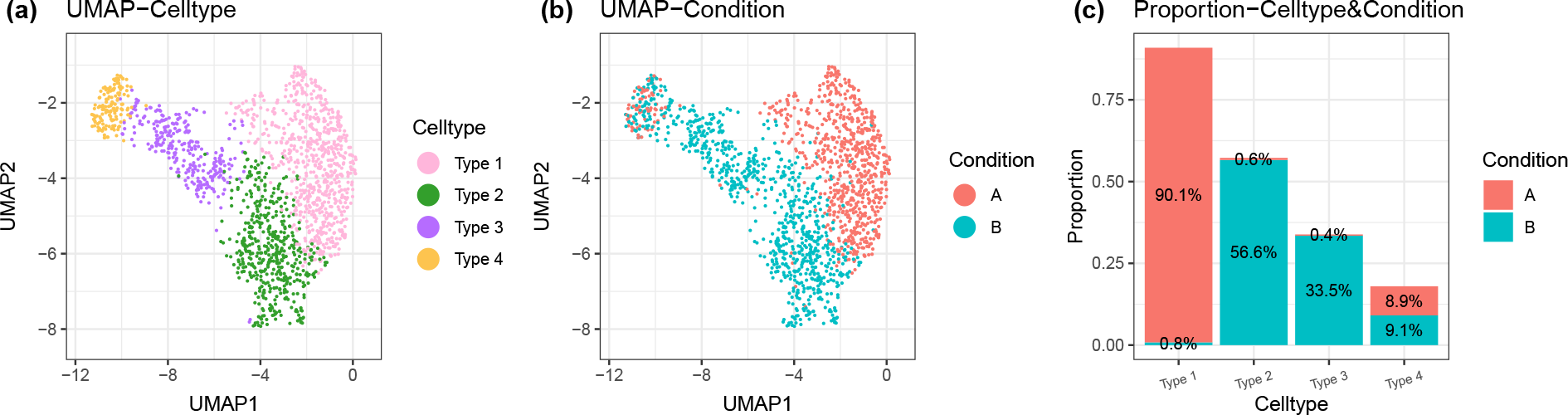
Simulation truth. UMAP plot of the single cell data colored by true cell type (panel a), and condition (b). Panel (c) shows the proportion of true cell types within the two conditions, respectively.

The inference results are consistent with the simulation truth. Figures 3 (a)-(c) show posterior summaries for the weights and the weight differences between the two conditions. Under the assumed model structure, the differences highlight which clusters are common and which are condition-specific. Figures 3 (c) and (d) indicate that cells of type 1 are mainly assigned to clusters specific to condition A, while cells of type 2 and 3 are mainly assigned to clusters specific to B, and type 4 cells are mostly assigned to the common clusters. Compared to the simulation truth that type 1 cells are enriched under A, type 2 and 3 cells are enriched under B, and type 4 cells are common, the accuracy of classifying cells to common, condition A enriched, and condition B enriched clusters is 91.2%. The balanced accuracy for the three types of clusters is summarized in Table 1.

**Table 1:**
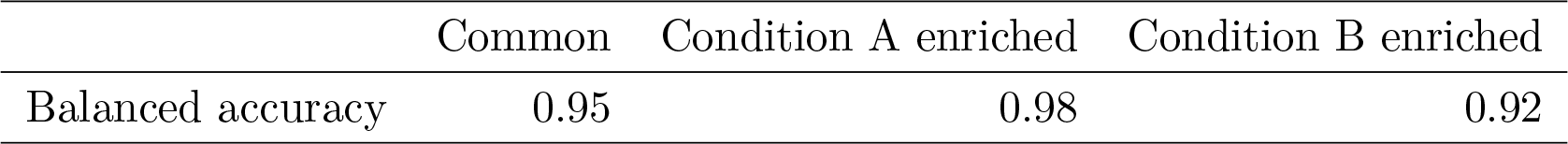
Balanced accuracy of classifying cells as Common/Condition A enriched/Condition B enriched.

**Figure 3:**
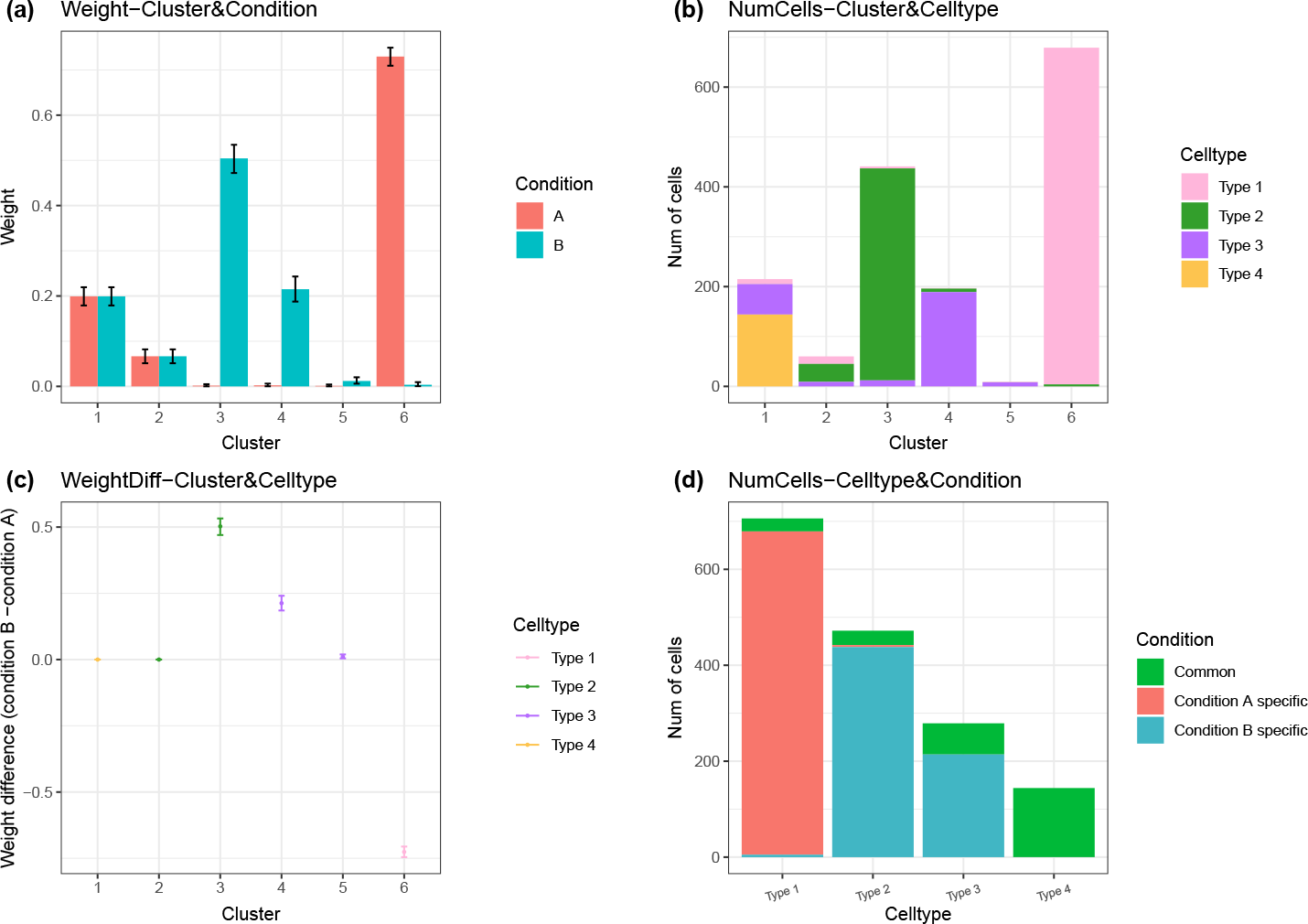
Inference results in the simulation study. (a) Posterior means and 95% credible intervals of the weights under the two conditions. (b) The number of cells from each true cell type in the reported clusters. (c) Posterior summaries of the differences for each cluster of the weights under condition A versus B. The plot shows the posterior means and credible intervals. (d) The number of cells in each true cell type being assigned into the three types of clusters.

Cluster assignments are shown in the panels in the first row of Figure 4. The figure plots UMAP scores colored by cluster assignments. The subfigures include a plot for all cells and for condition A and condition B only, respectively. Panels in rows 2 through 4 of Figure 4 show condition A cells and condition B cells in three types of clusters, common, specific to A, and specific to B. The clusters have the intended structures as desired. That is, the clusters specific to A have significantly higher proportion of cells from condition A, and very few cells from condition B, and vice versa for clusters specific to B.

**Figure 4:**
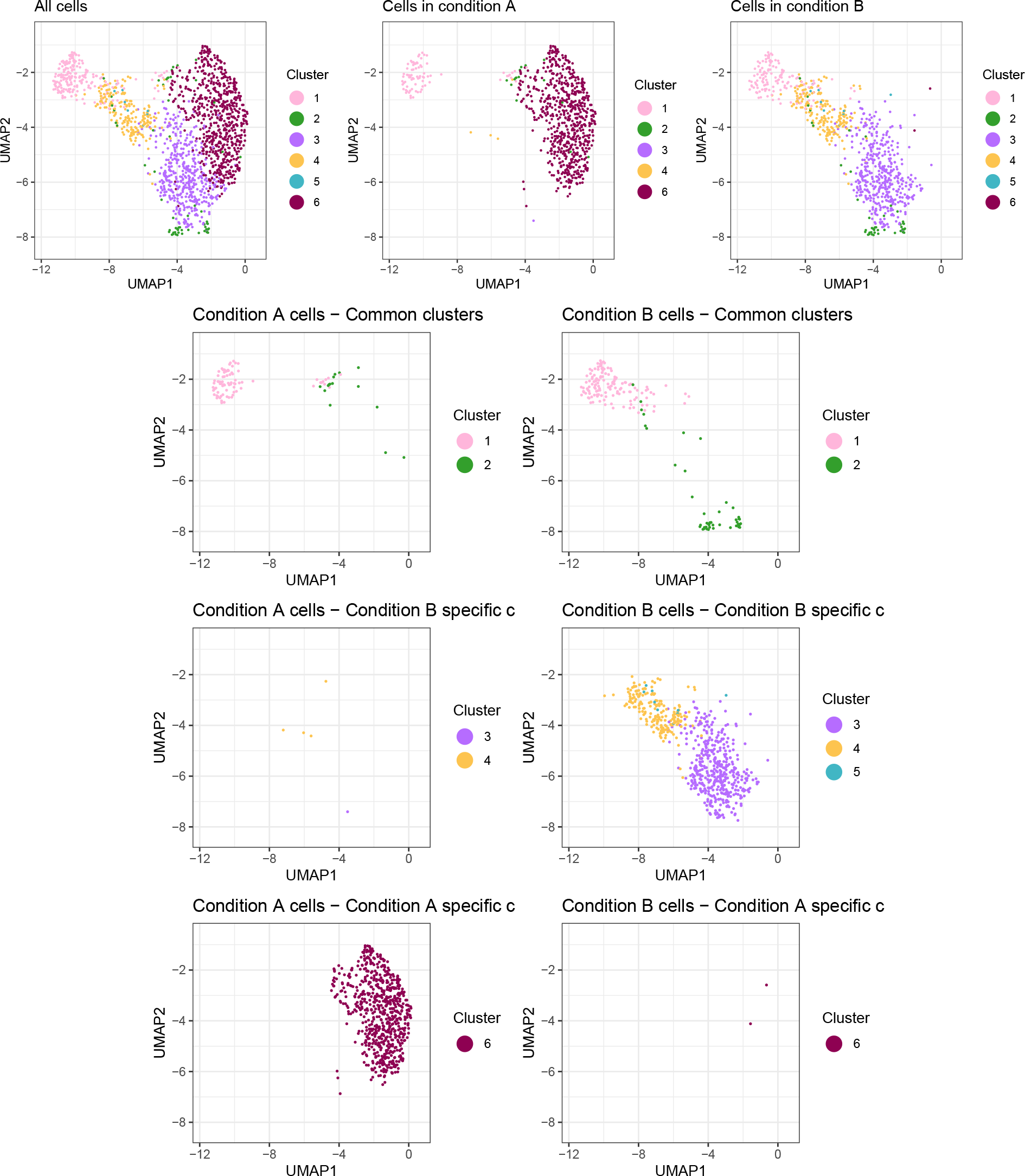
Simulation results (plotted as UMAP scores). The first row shows the clusters for all cells, cells under condition A, and cells under condition B, respectively. The second row shows condition A/B cells in the common clusters. The third row shows condition A/B cells in clusters that are enriched under condition B. The fourth row shows condition A/B cells in clusters enriched under condition A.

## 6 Results

We implemented inference under the 888 model on CRC data from Zhang et al. [22]. The features *y*_*ji*_ are the PCA scores obtained from the pre-processing pipeline outlined in Section 2. To prevent the PCA process from excluding representative immune cell biomarkers, we supplemented the features in *y*_*ji*_ with differentially expressed genes based on cell type annotation. Consequently, *y*_*ji*_ comprises the first *n*_1_ PCA scores and the top *n*_2_ differentially expressed (DE) genes based on false discovery rates. In our specific case, we choose the value of *n*_1_ based on an elbow plot of the proportion of variance explained versus the number of PCA components, suggesting *n*_1_ = 25, and we choose *n*_2_ = 20. We carry out separate inference for CD4+ and CD8+ cells due to the comprehensive research on the biomarkers specific to these cell types and the clear separation achieved through CD4+ and CD8+ annotation. In both, the CD4+ and CD8+ models, we set *H* to 2 as an upper bound on the resolution level of features that we can realistically expect in this data.

Posterior inference on the partitions reports the following homogeneous cell sub-populations. A point estimate of the partition, i.e., a clustering assignment, is obtained by minimizing variance-of-information (VI) loss [31]. The allocation of cells to these clusters under different conditions can be seen in Figure 6. Figures 5 (a) and (b) display the posterior means and credible intervals of the condition-specific weights and weight differences, respectively. Using the 888 model, we identified six clusters within the CD4+ T cell condition and five clusters within the CD8+ T cell condition. Specifically, within the CD4+ T cells, clusters 1, 2, and 3 were classified as common clusters, while clusters 4 and 5 exhibited enrichment in LOCRC. The (*H* + 1)-st common cluster here is an EO-specific cluster under the prior model for which we found *a posteriori* negligible difference of the condition-specific weights. Cluster 6, on the other hand, presented EO enrichment. Within CD8+ T cells, clusters 7 and 8 were classified as common clusters, cluster 9 showed LO enrichment, and Clusters 10 and 11 displayed EO enrichment. Another LO enriched CD8+ cluster has negligible weights under both conditions, and we combine it with the closest cluster.

**Figure 5:**
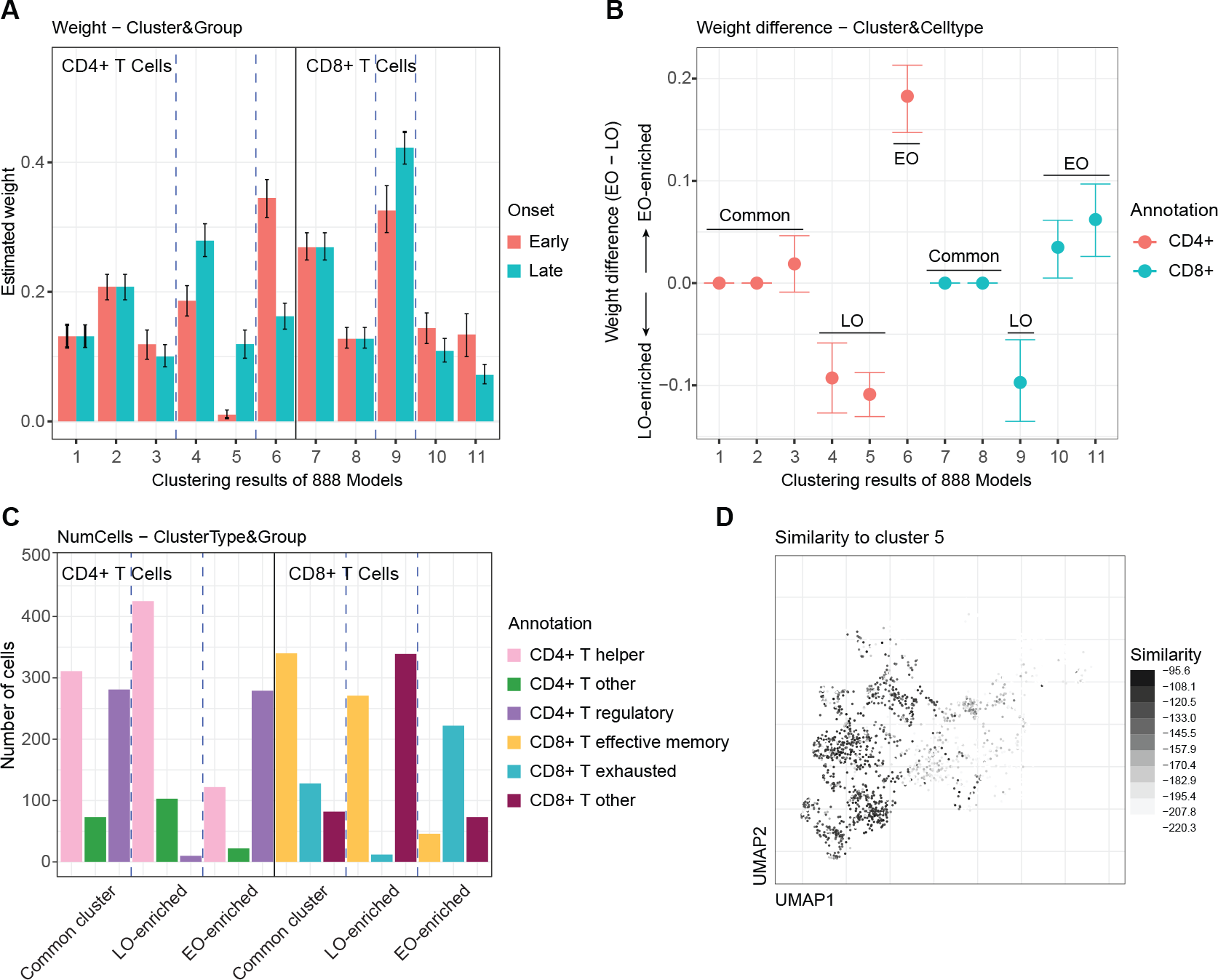
Results of the immune profile comparison between early-onset versus late-onset CRC. (a) Posterior mean and 95% credible interval of the cluster weights under EO/LO condition. (b) Posterior mean and 95% credible interval of the cluster weight differences colored by the dominant annotated cell type. (c) Number of cells from the annotated cell types in Common, LO-enriched, and EO-enriched clusters. (d) The similarity of each cell to cluster 5.

**Figure 6:**
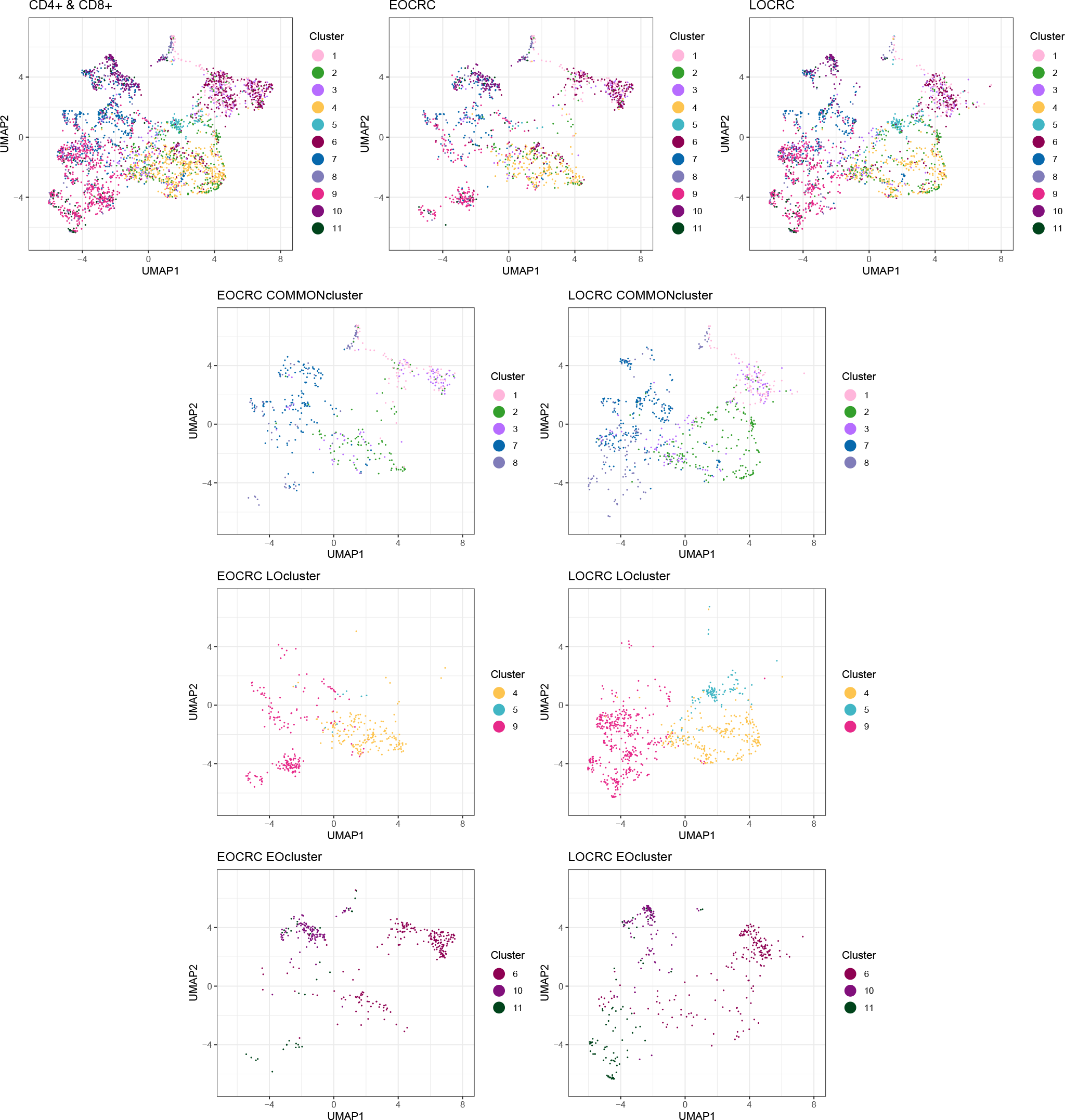
Results plotted in UMAP scores. The first row shows the clusters for all cells, EO cells, and LO cells. The second row shows EO/LO cells in the common clusters. The third row shows EO/LO cells in LO-enriched clusters. The fourth row shows EO/LO cells in EO-enriched clusters.

Next we studied the relation of the discovered clusters with known T cell subtypes, hoping to identify new variations or refined subtypes. Figure 5 (c) summarizes the composition of different immune cell types within the clusters. By comparing the 888 clustering results with the cell type annotation based on known markers, we made the following observations: (1) a specific subpopulation of regulatory T (Treg) cells demonstrated enrichment in the EOCRC samples; (2) a subpopulation of CD8+ effective memory T (Tem) cells displayed enrichment in the LOCRC samples; and (3) we identified a subpopulation of CD8+ exhausted T (Tex) cells that exhibited enrichment specifically in the LOCRC samples. Lastly, to illustrate the uncertainty in cluster assignment, Figure 5 (d) depicts the similarity of cells to a cluster, using cluster 5 as an example. This summary highlights the variability and ambiguity in cluster assignments.

Given the identified common, LO-enriched, and EO-enriched T cell subtypes, we look further into their gene expression profiles, hoping to annotate these clusters with known T cell states in cancer [33] (Figure 7). We identify some immune subtypes with differentially expressed genes matching the known markers. In the <u>LO-enriched CD4+ T helper</u> cells, we observed a high expression of CCR7 and TCF7, which are markers for naïve T cells, suggesting a self-renewal characteristic [34, 35]. Conversely, the <u>EO-enriched CD4+ T helper</u> cells exhibited high expression of CCL5, which was linked to various mechanisms of tumor progression, including extracellular matrix remodeling, enhanced tumor cell migration, and the induction of immunosuppressive polarization in macrophages [36]. In <u>EO-enriched CD4+ Treg</u> cells, we found higher expression of co-stimulatory molecules TNFRSF4 and TNFRSF18. These molecules are known to enhance the proliferation and activation of Treg cells [37]. Additionally, the <u>EO-enriched</u> Treg cells displayed higher expression of IL2RB, indicating potentially stronger immunosuppressive activity [38]. In the CD8+ T cell population, including Tem and Tex cells, the <u>EO-enriched Tem</u> cells exhibited higher expression of CXCL13. Previous studies have linked the presence of intratumoral CXCL13+ CD8+ T cells to unfavorable clinical outcomes and an immunoevasive environment. [39]. Additionally, the <u>EO-enriched Tex</u> cells showed higher expression of TNFRSF4, which contributes to immunosuppression and pro-angiogenesis in the tumor microenvironment [37].

**Figure 7:**
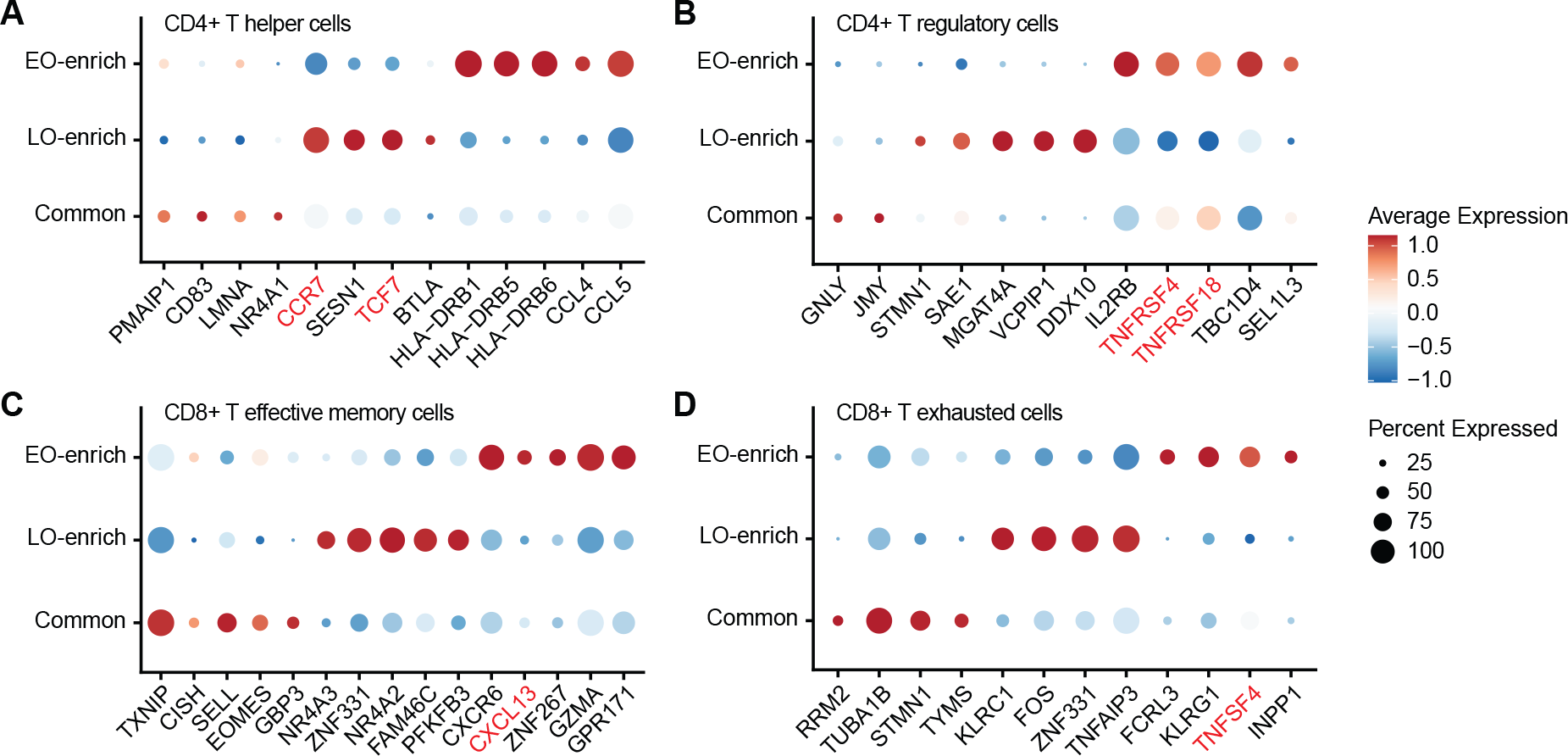
Differentially expressed genes in common, EO-enriched, LO-enriched clusters for CD4+ T helper, CD4+ T regulatory, CD8+ effective memory, and CD8+ T exhausted cells, respectively.

To summarize, we conducted comparative inference between two CRC conditions under the 888 model. Our findings revealed significant differences in immune cell subpopulations between EOCRC and LOCRC patients. The EOCRC samples, compared to LOCRC samples, exhibited an enrichment of immune-suppressive T cells, indicating a more immune-suppressive tumor microenvironment (TME) in EOCRC patients. These findings have the potential to inform novel approaches for the treatment and management of CRC, considering the distinct characteristics of the immune cell subpopulations in each condition.

## 7 Discussion

We introduced dependent mixture models with multi-layer structured priors on the weights to identify homogeneous subpopulations that highlight differential abundance across conditions. The proposed model allows us to evaluate comparative characteristics of T cell subtypes for EO versus LO colorectal cancer. We discussed limitations of existing models for dependent probability distributions in the context of the desired inference goals and showed how the asymmetric Dirichlet priors in the proposed hierarchically structured mixture models allow for the required comparison of immune profiles. Finite sample inference under the proposed approach was explored in simulation studies. Inference results with the immune profile comparison data, as intended, highlight immune subtypes that are enriched in EOCRC/LOCRC, and subtypes that are shared across the two conditions.

We only recommend full model-based inference using the proposed model when a quantitative comparison of subpopulations across conditions is desired. The proposed model is not needed and unnecessarily complex when the aim is only to report a cluster arrangement of experimental units – in that case any ad-hoc clustering algorithm would do.

Several extensions of the proposed model could be considered. In this implementation we used a fixed number of components *H* in the mixture. But the model could easily be extended to random *H*, with an informative prior. This generalization would give rise to additional computational challenges when implementing trans-dimensional posterior inference [19]. Another important generalization would be the inclusion of patient-specific random effects and possibly patient-specific covariates. One could introduce additive random effects for the normal location parameters in (7), or alternatively in the Dirichlet prior for the weights. In the latter case one would represent the Dirichlet random weights as normalized gamma random variables and use multiplicative random effects on those. The use of additive random effects formalizes the notion that the same homogeneous cell subpopulation might express slightly differently across patients or samples. The use of multiplicative random effects on the un-normalized weights corresponds to varying proportions of the same homogeneous subpopulations across patients. Incorporating covariates would naturally be formalized by multiplicative effects on the un-normalized weights.

The proposed model for comparative immune profiling might be used as a submodel for immune cells in a larger study of single cell data. Incorporating this submodel in the overall study pipeline, one can address the analysis on the immune subtypes that are distinctively enriched in some conditions, which helps explain the difference of micro-environment across disease subtypes.

## Appendix A

### Clinical characteristics of patients with CRC

**Table 2:** Clinical characteristics of 7 patients with CRC involved in this example.

| Characteristics | P0701 | P1012 | P1207 | P1212 | P1228 | P0215 | P0411 |
| --- | --- | --- | --- | --- | --- | --- | --- |
| Age | 68 | 35 | 66 | 42 | 77 | 75 | 75 |
| Gender | Female | Female | Female | Female | Female | Male | Male |
| Histological type<sup>a</sup> | Rectum<br>ADC | Colon<br>ADC | Colon<br>ADC | Colon<br>ADC | Colon<br>ADC | Colon<br>ADC | Rectum<br>ADC |
| pTNM: T | 1 | 4 | 4 | 4 | 4 | 4 | 4 |
| pTNM: N | 0 | 2 | 0 | 0 | 0 | 2 | 0 |
| pTNM: M | 0 | 0 | 0 | 0 | 0 | 1 | 0 |
| Stage | I | III C | II | II | II | IV | II B |
| Tumour size | 1×0.8 cm | 7×6 cm | 6×6 cm | 6×4 cm | 4.5×4 cm | 6.5×4 cm | 6.5×3.5 cm |
| Grade | Well-<br>differentiated | Low-<br>differentiated | Moderate-<br>differentiated | Low or moderate<br>differentiated | Low or moderate<br>differentiated | Low-<br>differentiated | Moderate-<br>differentiated |
| MSI status<sup>b</sup> | MSS | MSS | MSS | MSS | MSS | MSS | MSS |

<sup>a</sup> ADC, adenocarcinoma.
<sup>b</sup> MSS, microsatellite stability; MSI, microsatellite instability.

## Appendix B

### Proof of Proposition 1

**Proof:** We have introduced the cluster assignment indicators *s*_*ji*_, *c*_*ji*_ for unit *i* under condition *j*, where *c*_*ji*_ characterizes the type of cluster being common or certain condition-specific, *s*_*ji*_ is the cluster within a certain type. The probability function for the cluster assignment is given in equation (3). Assume we have a total number of *N* subjects under conditions *j* = 1, …, *J* . Let *𝒮* and *𝒞* denote the partition of [*N*] = *{*1, …, *N }* induced by *s*_*ji*_ and *c*_*ji*_, respectively. Suppose 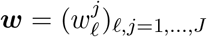 is a vector containing all the 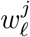 weight parameters. Marginalizing out *ϵ* and ***w***, we have

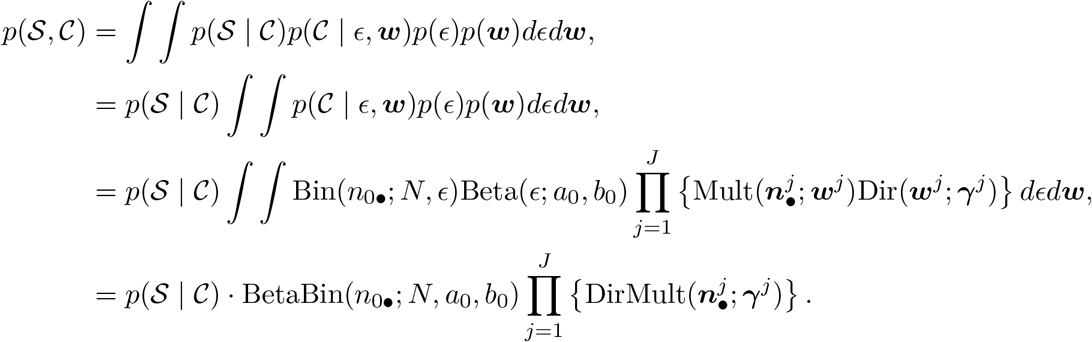

The second equation is true because *p*(*𝒮* | *𝒞*) is independent of *ϵ* and ***w***. The distribution of *p*(*𝒮* | *𝒞*) is a pEPPF induced by the EPPF of the MFMs 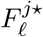 and *F*_0_. Each of them is given by equation (12), and the elements are conditionally independent given cluster type assignment *c*_*ji*_ = 0 or *ℓ*. Therefore, the pEPPF given the cluster type indicators *{c*_*ji*_*}* is

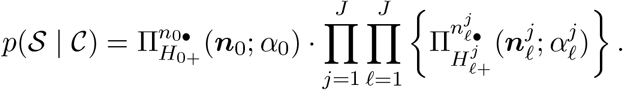

□

### Proof of Proposition 2

**Proof:** The Pólya-urn representation of the predictive probability function is derived from the random partition probability function for the model. The proof follows the notations in Section 4.

When a new unit under condition *j* is allocated to a new cluster in the set of common type clusters, such that 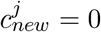, 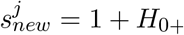, the predictive probability function is given by

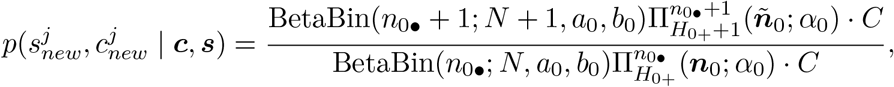

where 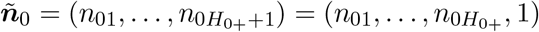, and

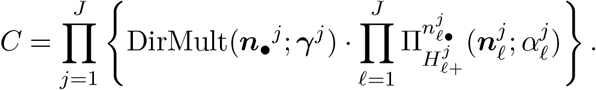

Substituting the beta-binomial probability function, the ratio of the *BetaBin*() factors is

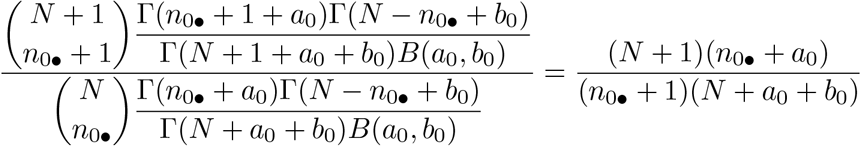

Plugging in the definition of Π(), the ratio of the Π() term is

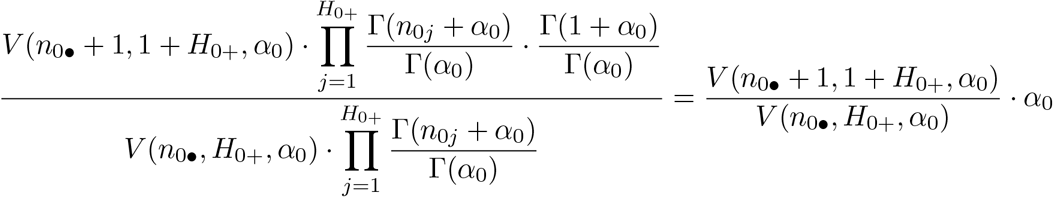

Therefore, when 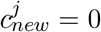, 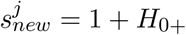,

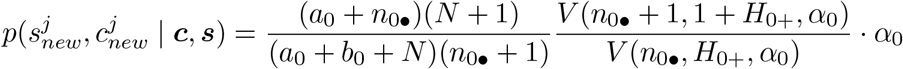

Similarly, when 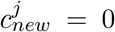, 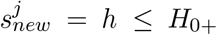, let ***ñ***_0_ = (*n*_01_, …, *n*_0*h*_ + 1, …, *n* 0*H*_0+_). The predictive probability function is

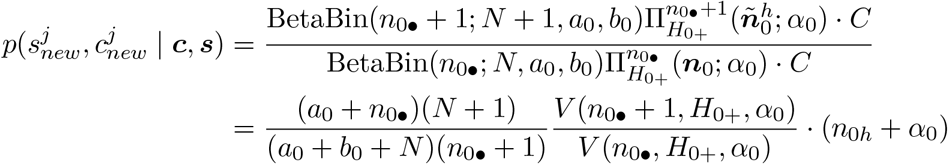

When 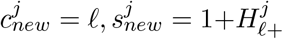 let 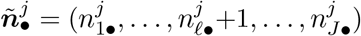 and 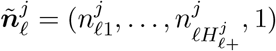.

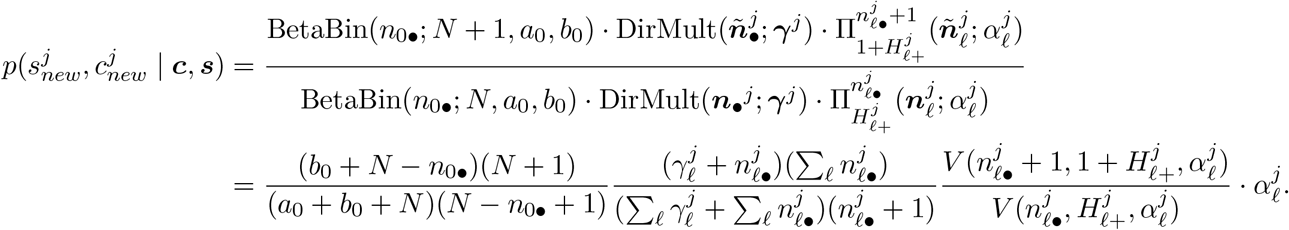

When 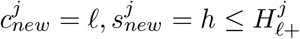, let 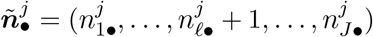 and 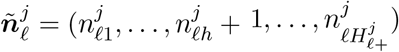.

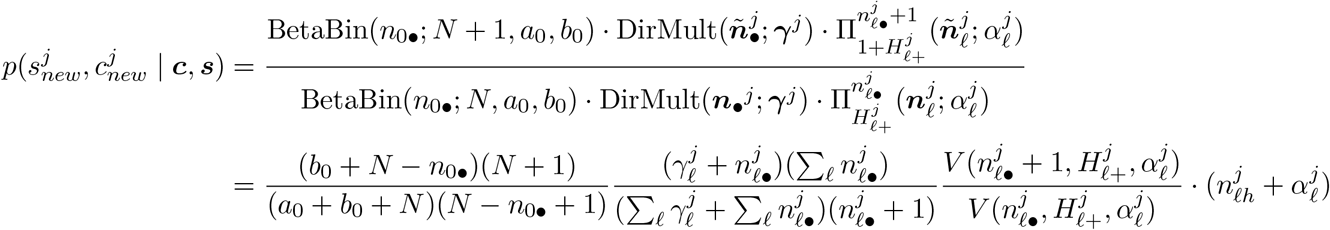

□

### Proof of Proposition 3

**Proof:** To prove Proposition 3 for any given *j ∈ {*1, …, *J }*, we consider the posterior contraction rate of the mixing measure *G*_*j*_ given observations 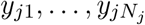. For the moment we drop all subscripts and superscripts *j* in the parameters. Then the 888 model under conditions (13) in Lemma 1 can be written as,

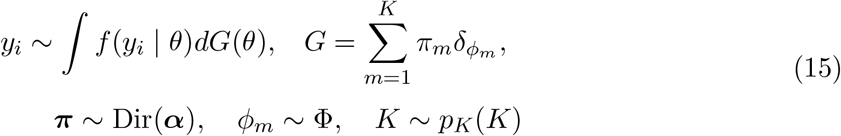

where 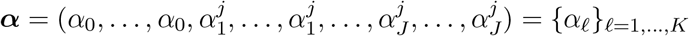.

Model (15) is a static MFM model if we replace the value of ***α*** by ***α*** = (*γ/K*, …, *γ/K*). Based on Guha et al. [29], under the following assumptions on the MFM model,

(P.1) The parameter space Θ is compact, while kernel density function *f* is first-order identifiable and admits the uniform Lipschitz property up to the first order.
(P.2) The base distribution Φ is absolutely continuous with respect to Lebesgue measure *μ* on ℝ and admits a density function *g*(*·*). Additionally, Φ is approximately uniform: min_*θ∈*Θ_ *g*(*θ*) *> c*_0_ *>* 0.
(P.3) There exists *ϵ*_0_ *>* 0 such that 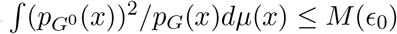 as long as *W*_1_(*G, G*^0^) *≤ ϵ*_0_ for any 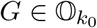 where *M* (*ϵ*_0_) depends only on *ϵ*_0_, *G*^0^, and Θ.
(P.4) The prior *p*_*K*_ places positive mass on the set of natural numbers, i.e., *p*_*K*_(*k*) *>* 0 for all *k ∈* ℕ.

we have that

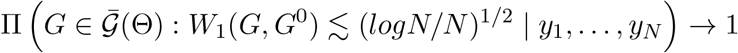

in probability under the true mixing measure *G*^0^.

We first prove that the model in equation (15) with asymmetric Dirichlet prior ***α***= {*α*_*ℓ*_}_*ℓ*=1,…,*K*_ has the same posterior contraction rate under conditions (P.1), (P.2), (P.3), and (P.4). Following the steps in the proof in Section 6 in [29], we claim that equation (18) in the paper is also true under the prior *Q* = Dir(***α***) when ***α*** is asymmetric, such that

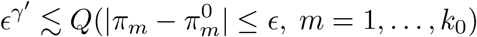

where *k*_0_ is the number of clusters in use, *γ*^*′*^ *>* 0 is a positive constant and *ϵ* is sufficiently small. Therefore, the conclusion on the contraction rate is proved by following the same steps as in Guha et al. [29]. We give the proof of the last claim:

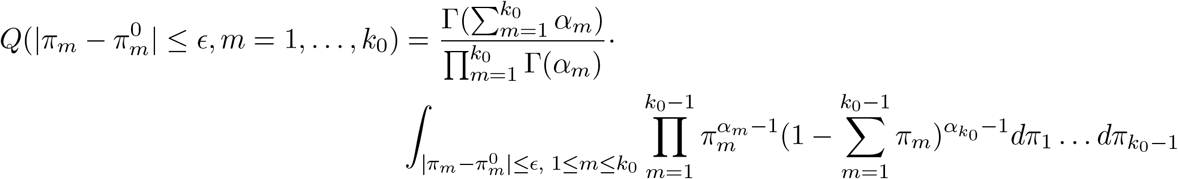

If 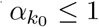, we have

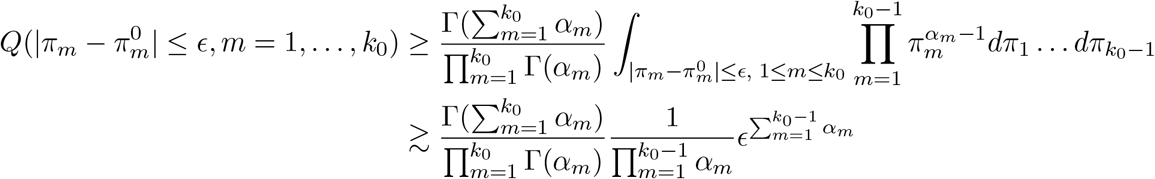

where the first inequality due to the fact that 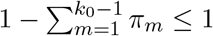.

If 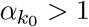, we have

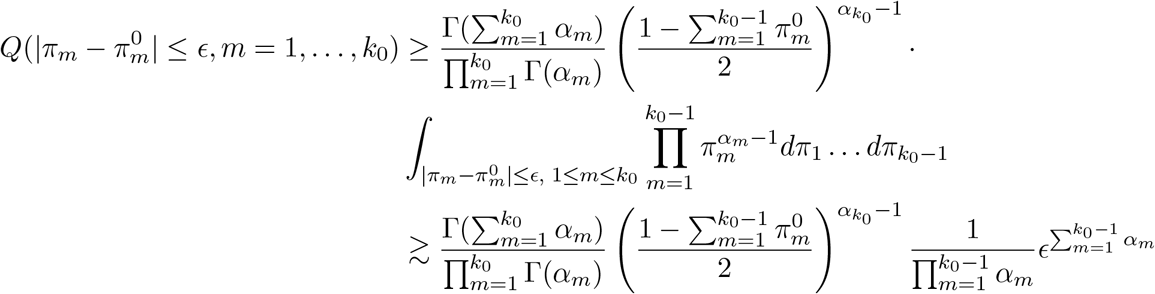

The first inequality in the above display is true because when 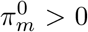 for all *m ∈ {*1, …, *k*_0_*}* for sufficient small *ϵ >* 0 such that 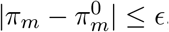, we have 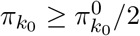.

We are now ready to show that the proposed 888 model satisfies the condition for consistency. We can restrict the parameters to a compact space, the kernel density *f* is a location-scale multivariate Gaussian distribution so that it is first order identifiable and admits the uniform Lipschitz property up to the first order (P.1), and the assumption (P.3) which controls the growing rate of KL neighborhood also holds for multivariate Gaussian. The prior distribution on the atoms in our model is continuous and approximately uniform (P.2). A prior *p*(*H*) for the number of clusters that places positive mass on the set of natural number is assumed so that (P.4) is satisfied. □

